# A bioengineered probiotic for the oral delivery of a peptide Kv1.3 channel blocker to treat rheumatoid arthritis

**DOI:** 10.1101/2022.07.12.499749

**Authors:** Yuqing Wang, Duolong Zhu, Laura C. Ortiz-Velez, J. Lance Perry, Michael W. Pennington, Joseph M. Hyser, Robert A. Britton, Christine Beeton

## Abstract

Engineered microbes for the delivery of biologics is a promising avenue for the treatment of various conditions such as chronic inflammatory disorders and metabolic disease. In this study, we developed a genetically engineered probiotic delivery system that delivers the small molecular biologic to the intestinal tract with high efficacy and minimized side effects. We constructed an inducible system in the probiotic *Lactobacillus reuteri* to secret functional Kv1.3 potassium blocker ShK-235 (LrS235). We show that LrS235 is capable of blocking Kv1.3 currents and preferentially inhibiting human T effector memory (T_EM_) cells proliferation *in vitro*. A single oral gavage of healthy rats with LrS235 resulted in adequate functional ShK-235 in the circulation to reduce inflammation in a delayed-type hypersensitivity model of atopic dermatitis mediated by T_EM_ cells. Furthermore, the daily oral gavage of LrS235 dramatically reduced clinical signs of disease and joint inflammation in rats with a model of rheumatoid arthritis without eliciting immunogenicity against ShK-235. This work demonstrates the efficacy of using probiotic *L. reuteri* as a novel oral delivery platform for the small molecule ShK-235, and provides a efficacious strategy to deliver other biologics with great translational potential.

**Significance Statement:** New therapeutics that combine efficacy with limited side effects and can be delivered non-invasively are needed to adequately treat patients with rheumatoid arthritis (RA) and other autoimmune diseases. Kv1.3 channel-expressing CCR7^-^ effector memory T (T_EM_) lymphocytes are significant players in the pathogenesis of multiple autoimmune diseases and blocking Kv1.3 reduces disease severity in rat models of RA and patients with plaque psoriasis. However, peptide therapeutics require repeated injections, reducing patient compliance. We used a bioengineered *Lactobacillus reuteri* as an oral delivery method of a Kv1.3 blocker for immunomodulation in rat models of atopic dermatitis and RA. This study demonstrates a novel approach for the non-invasive delivery of peptide-based therapeutics for the oral treatment of chronic inflammatory diseases.

## Introduction

Biologics now constitute a significant element of availbale medical treatments for various conditions such as chronic inflammatory disorders, cancer and metabolic disease. Nearly 30% of all drugs approved by the US. Food and Drug Administration in 2015-2018 were biologics (1), yet, the majority biologics are administered via parenteral route because of poor bioavailability via oral route. Fear of needles, injection-associated infection and pain are responsible for skipping doses by patients, especialiy for those with chronic inflammatory diseases that often requires life-long treatment. Rheumatoid arthritis (RA), one of the most common autoimmune diseases, mainly affects synovial joints but is also associated with a higher risk of cardiovascular, skeletal, psychological disorders and carries a significant socioeconomic burden (2). Although current therapeutics have considerably improved the management of RA in the last decade, RA-induced reduction in lifespan has not improved and maybe even worsened (3). Current RA treatments include nonsteroidal anti-inflammatory drugs (NSAIDs), corticosteroids, disease-modifying anti-rheumatic drugs (DMARDs), and biologic response modifiers such as antibodies targeting cytokines and their receptors (TNF-α or IL-1β, for example). These approaches focus on improving inflammation and pain control; however, they have little effect on the pathogenesis of RA and many increase the probability of infections or cancer. Many patients also experience significant side effects such as kidney or liver damage, leading 30–50% of patients to alter their treatment regimens (4, 5). Furthermore, the current biologics can be immunogenic and stimulate the generation of neutralizing antibodies after repeated drug administration (6, 7). Thus, new therapeutics that combine efficacy with limited side effects are needed to treat patients with RA adequately.

The pathogenesis role of T lymphocytes in RA has been extensively studied. Whereas CCR7^+^ naïve and central memory T (T_CM_) cells are the predominant T lymphocyte populations in the circulation and lymphoid organs, most T cells in the synovium and synovial fluid of patients with RA are CCR7^-^ effector memory T (T_EM_) cells, making them a desirable therapeutic target for RA (8, 9). At rest, human and rat T lymphocytes express low levels of two K^+^ channels, Kv1.3 and KCa3.1, that regulate plasma membrane potential and the homeostasis of Ca^2+^, a crucial second messenger in T cell activation (10-12). Upon activation, CCR7^-^ T_EM_ cells upregulate Kv1.3 while naïve and T_CM_ cells upregulate KCa3.1. Thus, T_EM_ cells are exquisitely sensitive to inhibition by Kv1.3 blockers. On the contrary, naïve and T_CM_ cells rely on KCa3.1 and escape Kv1.3 blockers. ShK-186 is a potent and selective peptide blocker of Kv1.3 that has been extensively tested in rats, non-human primates, healthy volunteers, and patients with a T_EM_ cell-mediated autoimmune disease in a Phase 1A/B clinical trial (10, 11, 13, 14). The *in vivo* safety and efficacy of ShK-186 were demonstrated, like for other biologics, after injections. The delivery of Kv1.3-blocking peptides via the buccal mucosa showed that a transmucosal route of delivery is feasible, albeit with low efficacy (15).

Here, we report the bioengineering of a probiotic for the oral delivery of ShK-235. ShK-235 is a recombinant analog of ShK-186, which cannot be produced recombinantly due to a non-amino acid adduct; we, therefore, generated ShK-235 that can be produced recombinantly and retains the potency and selectivity of ShK-186 for Kv1.3 channels (see supplemental Fig. S1A) (16). *Lactobacillus reuteri (L. reuteri)* as a species is an indigenous bacteria of the human and vertebrate animal gastrointestinal (GI) tract. It is one of the Lactic acid bacteria groups that has long been used as a cell factory in the food industry and is recognized as safe by the US Food and Drug Administration (17). It has an excellent safety profile in infants, children, adults, and even in an immunosuppressed population (18). The strain *L. reuteri* ATCC PTA 6475 is a well-characterized probiotic that does not colonize but survives transit through the GI tract of humans and rodents (19). It primarily resides in the proximal gastrointestinal tract but is also found in the urinary tract, skin, and breast milk (18). The benefits of *Lactobacillus reuteri* include modulating host immune response, enhancing gut mucosal integrity, inhibiting bacteria translocation, and promoting nutrient absorption (18). With the genetic tools being developed to facilitate the engineering of *L. reuteri* 6475 genomes, interest in using *L. reuteri* as a bioengineered tool for the oral delivery of therapeutics has increased.

In this project, we bioengineered *L. reuteri* to generate LrS235 that secretes ShK-235. The secreted peptide is functional in blocking Kv1.3 channels and suppressing the activation of T_EM_ lymphocytes *in vitro*. It crosses into the circulation after the oral gavage of healthy rats and displays good bioavailability and pharmacokinetics. Treatment with LrS235 effectively reduces disease severity in a rat model of RA, including joint inflammation and cartilage and bone damage, without immunogenicity.

## Results

### Bioengineering of *L. reuteri* LJ01 to secrete functional ShK-235

To generate a *L. reuteri* strain capable of secreting ShK-235, we codon-optimized the gene for ShK-235 for expression in *L. reuteri*, fused the usp45 signal peptide (20, 21) sequence to the 5’ end, and cloned it into the vector pSIP411 (22) resulting in plasmid pLL01. We have previously shown that usp45, a signal peptide identified from *Lactococcus lactis*, is capable of high level secretion of IL-22 from *L. reuteri* LJ01(21). pSIP411 allows for the controlled expression of genes by the addition of a peptide pheromone that activates a promoter, based on a two-component regulatory system identified in *L. sakei* (see Supplemental Fig. S1 for details). The resulting bacterial strain containing the pLL01 was named LrS235 (*L. reuteri* ShK-235). As a control we used pSIP411 contianing the *gusA* gene that encodes a β-glucoronidase (bacteria strain referred to as LrGusA thereafter). Induction of neither ShK-235 nor GusA secretion had any measurable effect on cells growth.

We assayed for the presence of active peptide by testing the ability of LrS235 supernatants to block Kv1.3 channels in mouse L929 fibroblast stably expressing mKv1.3 channels using whole-cell patch-clamp. LrS235 and LrGusA were grown to mid-exponential phase and induced for expression of ShK-235 or GusA and supernatants were processed as described in the materials and methods. As an additional control we used known concentrations of synthetic ShK-235 to plot a dose-response curve of the block of Kv1.3 (IC_50_ = 69 ± 24 pM; Supplemental Fig. S2). We next tested culture supernatants from LrS235 and LrGusA for Kv1.3 channel block and calculated the concentration of ShK-235 in supernatants using the dose-response curve. Supernatants from LrS235, but not those from LrGusA, blocked Kv1.3 currents (Fig. 1A). We calculated the mean concentration of ShK-235 in the supernatants from LrS235 to be 421.3 ± 75.9 pM (Fig. 1A, B and Supplemental Fig. S2. A, B), well above the IC_50_ of ShK-235 for Kv1.3.

**Figure 1.**
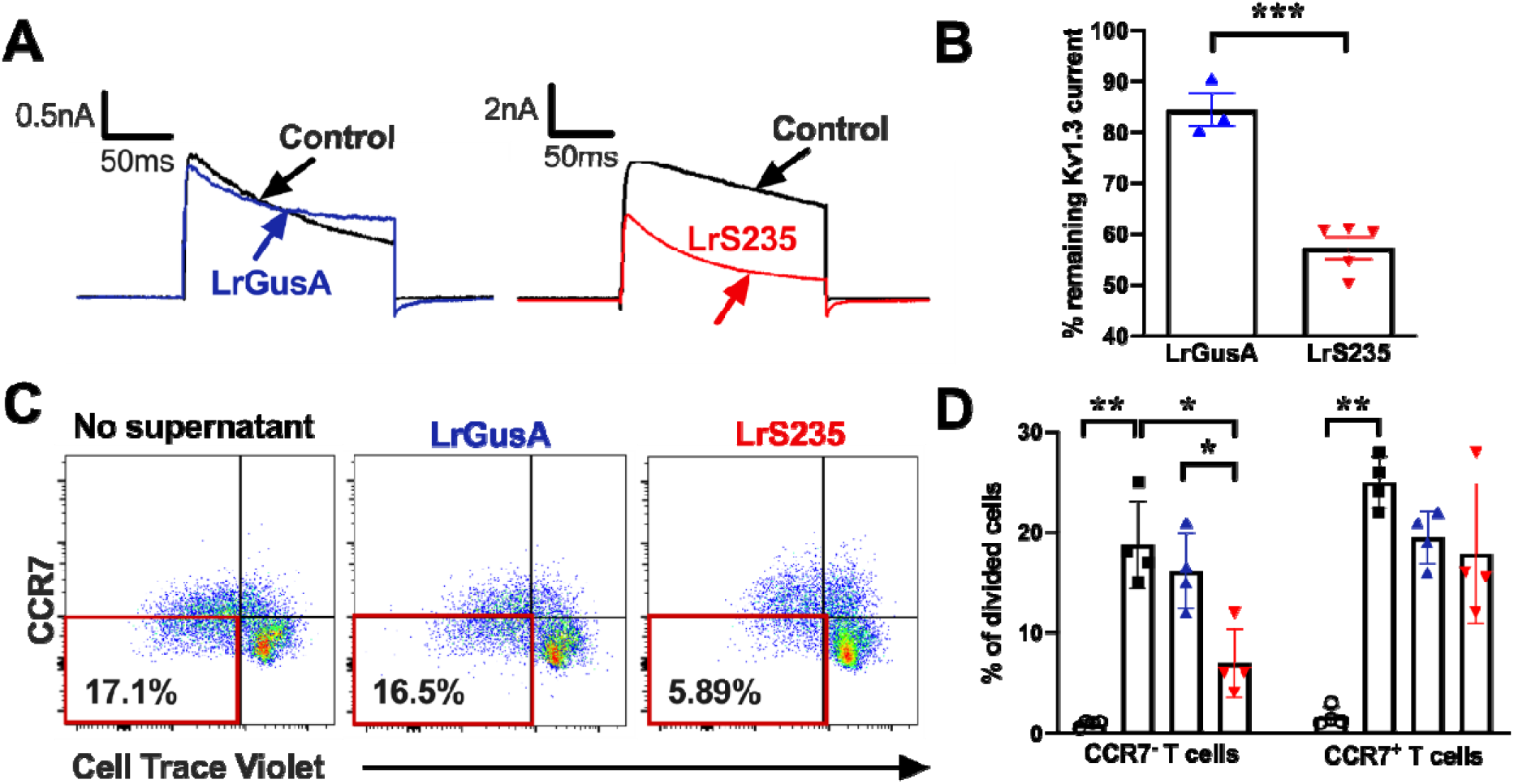
Supernatants from LrS235, but not from LrGusA, block Kv1.3 currents and inhibit the proliferation of human CCR7^-^ T_EM_ cells. (A) Representative whole-cell recordings of L929 cells stably expressing mKv1.3 before (control) and after the addition of supernatants diluted 1/10 of LrGusA or LrS235. (B) Percentage of remaining mKv1.3 currents after addition of LrGusA 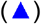 or LrS235 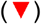 supernatants diluted 1/10. Mean ± SEM, each data point represents a different measurement. (C) Representative flow cytometry plots of CellTrace Violet dye dilution and CCR7 expression of CD3^+^ cells from human peripheral blood mononuclear cells stimulated for 7 days without any bacterial supernatant (left) or in the presence of supernatants from LrGusA (middle) or LrS235 (right). (D) Percent of divided human CCR7^-^ T_EM_ and CCR7^+^ naïve/T_CM_ cells in the absence of stimulation 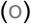 and after anti-CD3 induced stimulation in the presence of Lr medium (◼) or supernatants of LrGusA 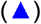 or LrS235 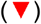 diluted 1/100. Mean ± SEM, N=4 different buffy coat donors. *p<0.05, **p<0.01.

To further validate LrS235’s production of active ShK-235, we tested the ability of LrS235 supernatants to inhibit the proliferation of T_EM_ cells. Incubation of supernatants from LrS235, but not from LrGusA, preferentially suppressed human CCR7^-^ T_EM_ cell proliferation by 63%, confirming the presence of biological active and physioloigically relevant ShK-235 peptide (Fig. 1C, D).

### Functional ShK-235 is detected in the circulation of healthy rats following the oral administration of LrS235

After validating that LrS235 produced and secreted functional ShK-235, we determined if oral gavage of the probiotic efficiently delivers functional ShK-235 to the circulation of rats. We first tested whether a compound of a molecular weight similar to that of ShK-235 can cross from the lumen of the GI tract into the circulation of healthy rats and rats with the collagen-induced arthritis (CIA) model of RA. The oral gavage of healthy rats or rats at the onset of CIA with 4 kDa dextran labeled with FITC showed that dextran could reach the circulation of both healthy and arthritic rats within 6 hours, with a higher permeability in the latter (Supplemental Fig. S3).

We orally administered a single dose of 10^9 colony-forming units (cfu) of LrS235 or LrGusA to healthy rats by oral gavage. Six hours later, we collected sera and tested the ability of serum samples to block Kv1.3 channels by whole-cell patch-clamp. Sera from rats given LrS235, diluted 1/100, inhibited Kv1.3 currents by 37%, whereas sera from rats administered with LrGusA had no effect on Kv1.3 currents (Fig. 2A, B). Based on the dose-response curve, the calculated concentration of LrS235 in the circulation 6 hours after a single oral gavage is 16 nM, significantly higher than the IC_50_ for Kv1.3 block of 69 ± 24 pM (Supplemental Fig. S2. A, B). As controls, we encapsulated 0.3 mg ShK-235 peptide or unflavored gelatin powder in the Torpac size 9h gelatin capsules, and enteric-coated these capsules with Acryl-EZE to target content delivery to the small intestine. Kv1.3 block was undetectable in both controls (Fig. 2A, B).

**Figure 2.**
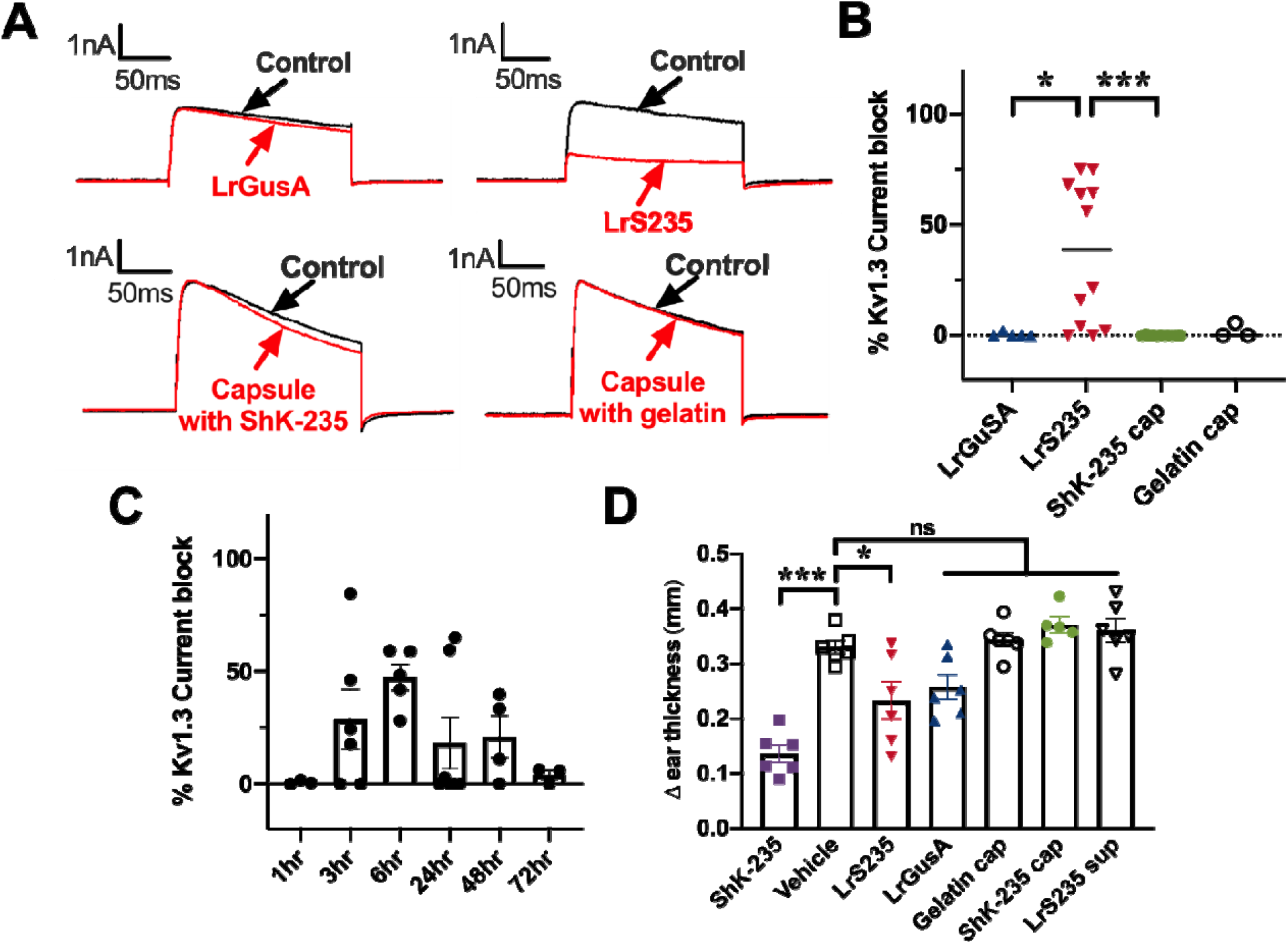
LrS235 secretes sufficient ShK-235 in the intestines for detection in the circulation of healthy rats. Healthy rats received an oral bolus of 1×10^9^ CFU of LrGusA 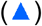 or LrS235 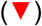, or an enteric-coated capsule filled with ShK-235 (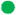, 2 mg/kg body weight) or gelatin (o). Blood was drawn at different time points and a single-cell patch-clamp was used to assess the ability of the serum to block Kv1.3 currents. N = 3 rats per time point. **A**, representative traces before (control) and after addition of serum diluted 1/100 from the 6-hour time point. **B**, Current block of serum samples collected at the 6-hour time point. N = 3 – 12 cells. **C**, Current block of serum samples collected at the indicated time points. each data point represents a different measurement. **D**, An active DTH reaction was induced against ovalbumin and rats received a single bolus of the following immediately before ear challenge: 1×10^9^ CFU of LrGusA 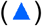 or LrS235 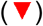 orally, an enteric-coated capsule filled with ShK-235 (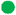, 2 mg/kg body weight) or gelatin (o) orally, 1 ml of LrS235 culture supernatant orally (⍰), or subcutaneous injection of 0.1 mg/kg synthetic ShK-235 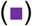 or vehicle (□). N = 6 rats per group (3 males, 3 females). **p<0.01, ***p<0.001.

We next determined the pharmacokinetics of ShK-235 after oral delivery of LrS235 by providing LrS235 by single oral gavage (10^9 cfu) to healthy rats and measuring blockage of Kv1.3 channels by whole cell patch clamp. Serum samples was collected at various timepoints ranging from 3 to 72 hours after the single bolus of LrS235. Peak levels of ShK-235 were detected 6 hours after LrS235 delivery, with the range of functional activity from 3 hours to 48 hours postgavage (Fig. 2C). These results demonstrate bacteria-delivered peptides to the gut can enter the serum and remain functional.

We then assessed if LrS235 gavage could reduce a delayed-type hypersensitivity (DTH) reaction in the ear of rats, a local autoinflammation mediated by antigen-specific T_EM_ lymphocytes (23-25). As expected, the oral gavage of LrS235 and the injection of ShK-235 significantly reduced inflammation by 30% and 58%, respectively, compared with the vehicle control (Fig. 2D). In contrast, treatment with LrGusA, capsules filled with ShK-235 or gelatin, and supernatants from the culture of the LrS235 had no significant effects on inflammation.

### LrS235 administration stops disease progression and bone and joint damage in collagen-induced arthritis in rats

Since we detected functional ShK-235 *in vitro* and in the circulation of rats, we sought to assess its efficacy in a model of arthritis in Lewis rats induced by porcine collegen II. Four groups of rats were treated with P6N buffer vehicle or synthetic ShK-235 injections, or LrS235 or LrGusA by oral gavage, starting from the onset of clinical signs. All vehicle-treated animals developed severe arthritis with a mean score of 26 ± 3 (Fig. 3A). The administration of LrGusA did not affect overall disease severity (mean score 25 ± 2). In contrast, the injection of synthetic ShK-235 reduced the mean score by ∼60% to 11 ± 3. The administration of LrS235 was even more effective with a mean score of only 4 ± 1, or an 84% reduction when compared to the vehicle control group. Histology performed on joints collected at the end of the *in vivo* trials showed severe cartilage degradation and erosion, angiogenesis, pannus formation, and synovial hyperplasia and inflammation in the CIA control rats (Fig. 3B, C). Whereas LrGusA had no benefits in any of these parameters, both synthetic ShK-235 and LrS235 significantly reduced all parameters, and no significant differences were seen between healthy controls and ShK-235 or LrS235 treated groups (Supplemental Table S1). Micro-CT imaging of hind limbs shows severe bone erosions in the CIA rats treated with vehicle or LrGusA, and better-preserved bones in CIA rats treated with synthetic ShK-235 or LrS235 (Fig. 3D). The zoomed, axial and pseudocoloral images are showed in Supplemental Fig. S4 and in the video of Supplemental Fig. S5.

**Figure 3.**
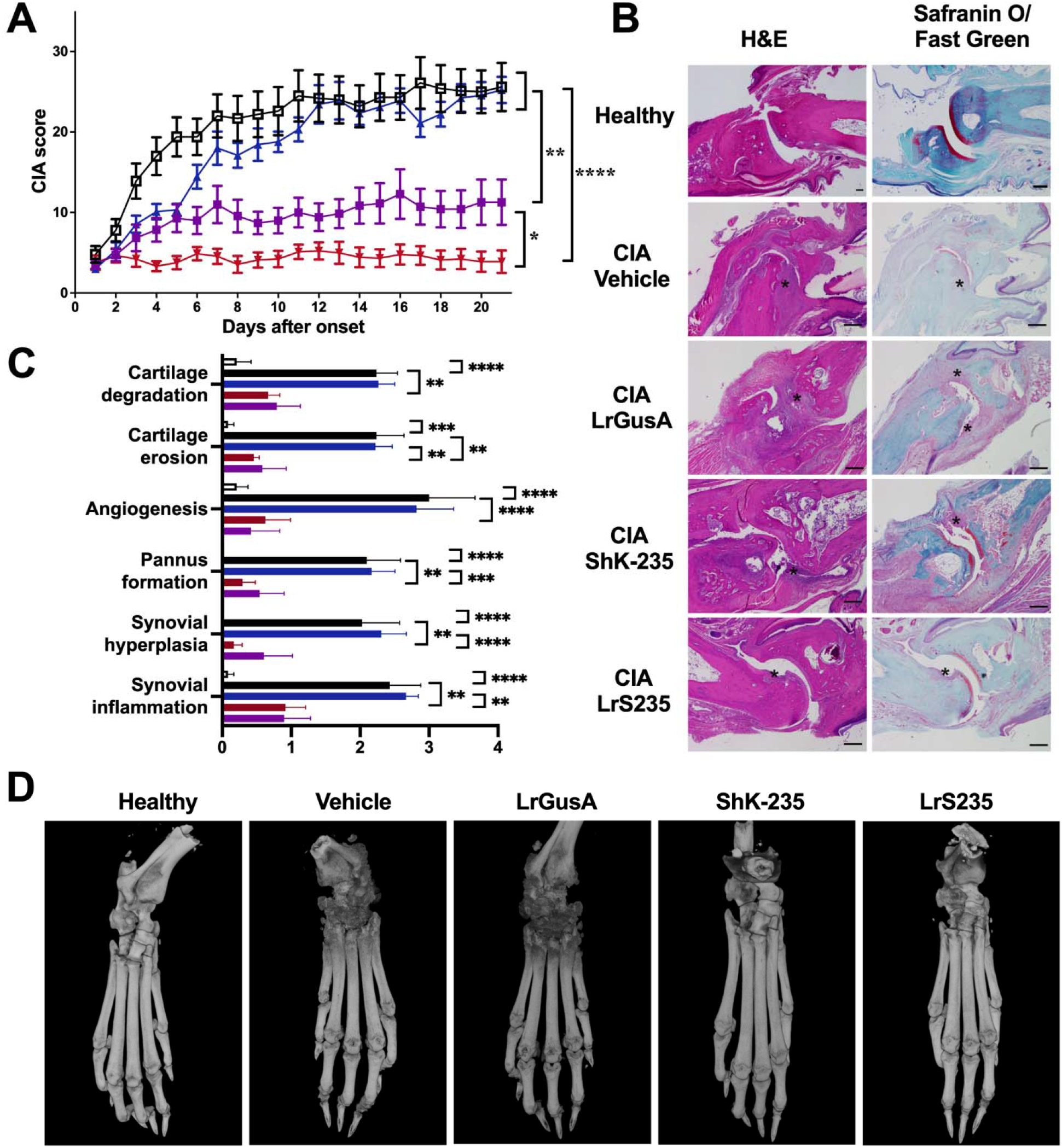
LrS235 stops disease progression, reduces bone and joint damage and inflammation in rats with collagen-induced arthritis. **A**. Clinical scores of paw inflammation from rats with CIA treated with vehicle (□) or with 100 μg/kg ShK-235 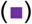 every other day starting disease onset, 1×10^9^ CFU LrGusA 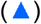 or LrS235 gavage daily 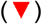. **B**. Hematoxylin & eosin (left) and safranin O/fast green (right) staining and histology scoring **(C)** of joints from paws from CIA rats received different treatments. Original magnification, 10x, scale bars, 100 μm. **D**. Representative micro-CT of paws from CIA rats treated with vehicle, synthetic ShK-235 every other day, or oral gavage with LrGusA, LrS235 daily. Data presented as mean ± SEM. N = 7 to 10 rats per group. Asterisks indicate areas of cartilage erosions. *P < 0.05; **P < 0.01, ***P< 0.001, ****P< 0.0001.

### ShK-235 produced by LrS235 is not immunogenic

To test if ShK-235 produced by LrS235 is immunogenic, we assessed the sera of rats at the end of the 21-day CIA trial shown in Fig. 3 for anti-ShK-235 IgG by ELISA assay. Plates coated with collagen II and HsTX1[R14A] were used as controls. HsTX1[R14A] is another peptide blocker of Kv1.3 with no sequence or structural homology to ShK-235. As expected, we detected high titers of antibodies against collagen II in rats with CIA, but only low titers of anti-ShK-235 antibodies (Fig. 4A and Supplemental Fig. S5). Similar low reactivity was observed against HsTX1[R14A], suggesting that the signal detected against ShK-235 was non-specific. We also performed a neutralization assay with the same CIA rat serum samples, and no neutralization of ShK-235 was detected, showing the absence of ShK-235 neutralizing antibodies (Fig. 4B).

**Figure 4.**
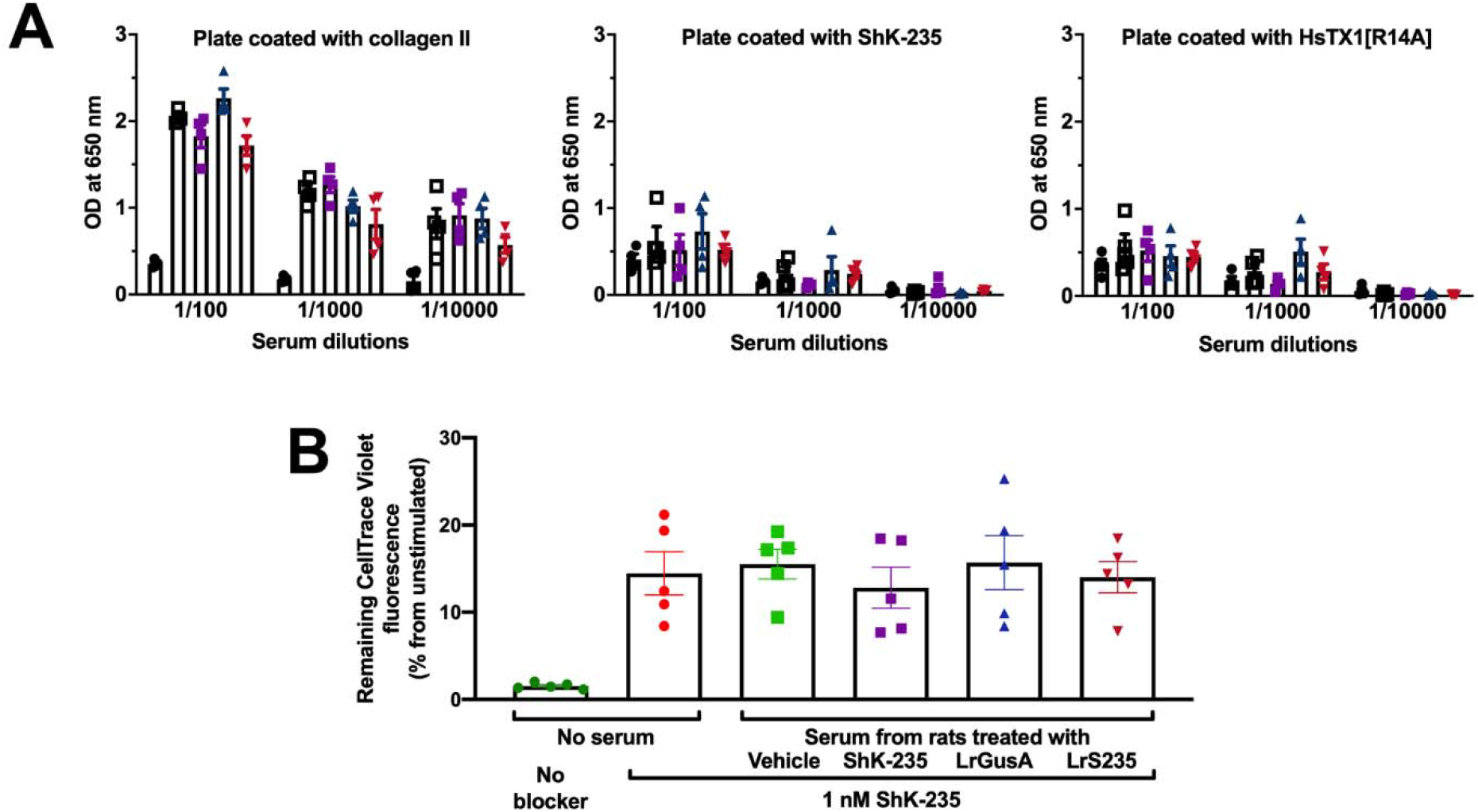
ShK-235, delivered via injection or LrS235, is not immunogenic. **A**. ELISA plates were coated with 10 μg/ml of either porcine collagen II, ShK-235, or HsTX1[R14A]. Sera from rats with CIA treated with vehicle, ShK-235, LrGusA, or LrS235 were tested at 10-fold dilution steps starting from a 1:100 dilution. The serum from non-immunized rats was used as a negative control. Each column represents the mean OD of 4 individual rat sera ± SEM. **B**. Data from different PBMC donors were normalized to the unstimulated cells with 100% CellTrace violet. The higher dilution of CellTrace Violet indicates a higher division rate. Data are present as Mean ± SEM.

To further investigate if long-term exposure of rats to LrS235 would elicit antibodies against ShK-235, we treated healthy rats daily with LrS235 or LrGusA for 8 weeks, stopped treatment for 12 weeks, followed by additional daily gavages of either LrS235 or LrGusA for 1 week. ELISA performed on blood samples collected either at the 8 week or the 21 week timepoints showed no IgG with specificity for ShK-235 was produced (Supplemental Fig. S7 A, B). This is expected as in the previous phase 1b trial for ShK-186, none of the paitents received ShK-186 by subcutaneous injection developed anti-peptide antibodies (14).

## Discussion

We have developed a bioengineered probiotic, LrS235, that secretes functional ShK-235 both *in vitro* and *in vivo*. Our engineering strategy enabled the production of an inducible peptide secretion system that fused ShK-235 to the usp45 secretion peptide, secretion of which can be induced upon the addition of the SppIP peptide pheromone. Oral gavage of the LrS235 daily starting at the onset of CIA in Lewis rats results in a reduction in disease severity. The protective effects of LrS235 in CIA are accompanied by a reduction in bone and joint damage with no immunogenicity.

Most peptide-based drugs require parenteral administration due to poor oral bioavailability (26). Those biologics are usually used for chronic diseases, and repetitive injections during the long-term disease course could reduce patient compliance and lead to poor outcomes. The convenience and widespread use of oral administration makes it the most desirable method in a clinical setting. Engineered probiotics can be used to deliver drugs or enzymes to treat metabolic disorders into the GI tract or tumors (27, 28). A genetically engineered strain of *E. coli* Nissle was used to deliver the angiogenesis inhibitor tumstatin to treat murine melanoma (29). More recently, the Phe-metabolizing enzyme phenylalanine ammonia lyase (PAL) and L-amino acid deaminase (LAAD) enzyme expressing *E*.*coli* Nissle has completed human phase 1/2a clinical trials and yielded positive results in phenylketonuria patients (30), demonstrating the potential of using bacteria as a promising drug delivery method. However, in those successful experiments, tumstatin was delivered via intrapetitoneal injection of the bioengineered bacetria that were subsequently cleared by the immune system and LAAD was effective directly in the GI tract. Here, we show that the engineered probiotic LrS235 can be given orally with a good bioavailability of the ShK-235 it produces in the circulation of the rats. In contrast, the oral administration of a high dose of synthetic ShK-235 packaged into enteric-coated capsules yielded no detectable ShK-235 in the circulation of the rats and no benefits in the DTH model, and neither did the oral gavage of supernatants from the culture of LrS235 affect DTH. This is consistent with prior work showing low oral bioavailability of peptide and protein therapeutics due to the hydrophilic properties of the peptides and the presence of potent peptidases and proteases in the GI tract (26). The bioavailability of venom-derived Kv1.3-blocking peptides through mucosal membranes has been demonstrated after buccal and pulmonary delivery (15, 31), suggesting the feasibility of delivery via the intestinal mucosa. A probiotic-based delivery presents several advantages as the daily ingestion of probiotics by humans is well established, and *L. reuteri* 6475 has already been tested in humans for research purposes and is also available as a commercially available supplement (Osfortis™). In addition, *L. reuteri* will continuously deliver ShK-235 throughout the GI tract, whereas delivery via capsule is pH-dependent and thus restricted to a one-time bolus to the distal intestine. Furthermore, *L. reuteri* has an affinity to mucus and will deliver the peptide in close proximity to the intestinal epithelium. Many bacteria produce extracellular vesicles that deliver bacterial products to the host’s circulation or tissues (32). More work is needed to determine whether such vesicles are involved in the remarkable bioavailability of ShK-235 following the oral delivery of LrS235.

LrS235 had significantly higher efficacy than the injected synthetic ShK-235 in reducing disease severity in CIA rats. At first glance, this could be explained by the administration frequency, LrS235 being given daily while the synthetic peptide was injected every other day. However, we had tested Kv1.3 blocker administration at frequencies ranging from every 6 hours to once a week and found that injections every other day provided a peak efficacy not improved by more frequent injections. This is likely due to the natural depot of the peptide at the site of subcutaneous injection, which leads to a slow release in the circulation (14). Another likely explanation is that the probiotic itself has anti-inflammatory effects. *L. reuteri* decreases intestinal inflammation in several mouse models (19, 33). In our CIA trials, L. reuteri producing GusA instead of ShK-235 reduced disease severity at the early stage of CIA, although significantly only on days 3 and 5 after the onset of clinical signs. Since rheumatoid arthritis is a chronic inflammatory disease, the probiotic and ShK-235 may synergistically suppress inflammation and improve the therapeutic effects of LrS235. While Kv1.3 blockers preferentially target T_EM_ lymphocytes, *L. reuteri* products may target effectors of the innate immune system, such as macrophages and dendritic cells, thus showing a modest protective effect early in CIA progression. Since CIA induction involves incomplete Freund’s adjuvant that induces a strong inflammation, LrGusA may show more anti-inflammatory benefits in milder models of inflammation. In RA and its animal models, joint inflammation disturbs the balance between osteoclasts and osteoblasts by inhibiting osteoblast differentiation and augmenting osteoclast function (34). Indeed, probiotics have been shown to regulate bone health through secretion of peptides or lipids, modulation of the host immune system, modifying the gut microbiome, and influencing local pH (35-37). Finally, the intestinal microbiome is perturbed in RA and its animal models (38), it is possible that the rescue of dysbiosis by LrS235 treatment accounts for the improved efficacy than synthetic ShK-235 injections. Further work is needed to validate this hypothesis.

Prior studies with injected blockers of the Kv1.3 channels, including ShK analogs, showed that a high efficacy in reducing the severity of CIA was accompanied by a reduction in DTH severity. Whereas both injected synthetic ShK-235 and ShK-235 delivered orally via LrS235 reduced inflammation in both CIA and DTH, LrS235 was more effective in CIA, and synthetic ShK-235 was more effective in DTH. Following subcutaneous injection, ShK and its analogs reach the circulation in less than 30 min (24). In contrast, the peak concentration of ShK-235 in the circulation of healthy rats is achieved 6 hours after a single dose oral delivery of LrS235. After the DTH challenge with ovalbumin, ovalbumin-specific T_EM_ cells enter the site of challenge and interact with local antigen-presenting cells within 3 hours (23). We administered both synthetic ShK-235 and LrS235 at the time of challenge, giving sufficient time for the synthetic peptide to reach the circulation and the T_EM_ cells before the start of the local inflammatory response but inflammation at the site of challenge had already begun by the time ShK-235 produced by LrS235 reached the circulation. Furthermore, other charged peptides with structures similar to that of ShK-235 accumulate in the joints following systemic administration (39, 40). This unique, and not fully understood feature of these peptides likely also plays a role in the remarkable efficacy of ShK-235 in reducing disease severity in CIA.

Many drugs, and especially biologics, are immunogenic and the resulting neutralizing antibodies can eventually render the medications ineffective. ShK is highly homologous to a domain of MMP-23 (41) and thus resembles a self-peptide, and is likely recognized as such by the immune system. As a result, ShK and its analogs induce little to no immunogenicity in rats or humans (9, 14) and we have found no immunogenicity of ShK-235 delivered via LrS235 in either healthy or CIA rats, regardless of short or long-term treatment.

In our immunogenicity ELISA, we used porcine collagen II as a positive control for detecting IgG as the rats had been immunized against this protein to induce CIA. There was no difference in anti-porcine collagen II IgG titers between the rats in the different treatment groups, this suggests that the B lymphocytes producing these antibodies belong to the CD27^-^IgD^-^ memory subset that relies on KCa3.1 rather than Kv1.3 channels for their function (42). The frequency of CD27^-^IgD^-^ B cells is increased in patients with both early and established RA (43) and is likely the subset of B cells producing the IgGs detected in the rats with CIA.

Altogether, we have developed the engineered ShK-235 secreting probiotic LrS235 that treats the animal model of rheumatoid arthritis efficiently. Our findings provide an alternative delivery strategy for peptide-based drugs and suggest that such techniques and principles can be applied to a broader range of drugs and the treatment of chronic inflammatory diseases.

## Materials and Methods

### Construct design for the Inducible Expression of ShK-235

ShK-235 differs from ShK by a Q16K substitution, an I21M substitution, and the addition of an Ala to the C-terminus (Supplemental Figure S1A). Codon optimized ShK-235 with signal peptide Usp45 was synthesized and ligated into NcoI-EcoRI digested pSIP411 generated pLL01 (Supplemental Figure S1B). The ShK-235 seceration plasmid pLL01 was enlarged in E. coli 1000 and was then electransformed into L. reuteri 647 competent cell resulting in ShK expression strain LrS235. All LrS235 or LrGusA used in this paper were induced with induction peptide unless otherwise stated.

### Synthesis of ShK-235 and HsTX1[R14A]

Both Kv1.3-blocking peptides were previously described (16, 44). ShK-235 and HsTX1[R14A] were synthesized using an Fmoc-tBu solid-phase synthesis strategy. Each coupling was mediated with diisopropyl carbodimide in the presence of HOBT. All deprotections were accomplished with 20% piperidine in dimethyl formamide. Following synthesis of the linear chain, each peptide was cleaved and deprotected using trifluoroacetic acid (90%) with carbocation scavengers (triisopropyl silane, H_2_O, DODT and thioanisole, (2% of each v/v)) for 3 hrs at ambient temperature. The peptides were each precipitated into methyl t-butyl ether. The linear peptides were purified by RP-HPLC and subsequently oxidatively folded in the presence of glutathione in ammonium acetate buffered aqueous solution. The cyclized products were isolated by RP-HPLC and fractions with a purity >95% by analytical HPLC were subsequently pooled and lyophilized. Each peptide was found to have expected theoretic mass for the formation of 3 and 4 disulfide bonds, respectively, for ShK-235 and HsTX1[R14A]. Synthetic ShK-235 and HsTX1[R14A] were dissolved in P6N buffer (10□mM sodium phosphate, 0.8% NaCl, 0.05% polysorbate 20, pH□6.0) to prepare stocks at 1 mg/ml (14, 24).

### Animals

Male and female Lewis rats (7-8□weeks old; Envigo, Indianapolis, IN, USA) were group-housed and provided food and water *ad libitum*. All animals were housed in a facility accredited by the Association for Assessment and Accreditation of Laboratory Animal Care International-accredited. All experiments involving rats were approved by the Institutional Animal Care and Use Committee at Baylor College of Medicine.

### Patch-clamp electrophysiology

To determine the concentration of ShK-235 in the LrS235 supernatants and rat sera, a dose-response of ShK-235 block of Kv1.3 was determined by adding known concentrations of the peptide to naïve rat serum and then testing by whole-cell patch-clamp of mouse L929 fibroblasts stably expressing mKv1.3 channels (45) using a Port-a-Patch automated patch-clamp system (Nanion, Livingston, NJ), as described (16, 46). Culture supernatants and serum samples were then assessed with the same technique, and the dose-response curve was used to determine peptide concentration.

### Human T lymphocyte proliferation assays

Buffy coats were purchased from the Gulf Coast Regional Blood Center (Houston, TX), and mononuclear cells were enriched using Histopaque-1077. The cells were loaded with 5 µM CellTrace Violet (Invitrogen, Waltham, MA) according to the manufacturer’s instructions (47) and incubated for 30 min with sterile-filtered supernatants from LrS235 or LrGusA, buffered to pH 7.4 and diluted 1/10 in tissue culture media, before the addition of anti-CD3 antibodies (clone OKT3, 1 ng/ml, 037-85, Thermo Fisher, San Francisco, CA). Seven days later, cells were stained with anti-CD3 antibodies conjugated to phycoerythrin (BioLegend 300308, lot B209105) and anti-CCR7 antibodies conjugated to FITC (R&D Systems, FAB197F, lot LEU1615081) and the dilution of CellTrace Violet in CD3^+^CCR7^-^ T_EM_ cells and CD3^+^CCR7^+^ naive/T_CM_ cells was measured by flow cytometry on a BD CantoII as quantification of cell proliferation. Data were analyzed with FlowJo.

### Encapsulation of ShK-235

Torpac size 9h gelatin capsules (Fairfield, NJ) were filled with either 0.3 mg synthetic ShK-235 or unflavored gelatin powder (Kraft Heinz, Chicago, IL) using the Torpac dosing kit. Each capsule was enteric-coated with Acryl-EZE® (Colorcon, Harleysville, PA) prior to delivery via oral gavage to target content delivery to the small intestine.

### Bioavailability and pharmacokinetics of ShK-235 after oral delivery

Male rats received a single oral gavage of either 1×10^9 cfus LrS235 or an enteric-coated capsule filled with ShK-235. Blood was collected via the saphenous vein at the different time points indicated in the figure, the last blood draw being a terminal cardiac puncture (48). Serum samples were assayed for Kv1.3 block by patch-clamp electrophysiology.

### Induction and monitoring of an active delayed-type hypersensitivity reaction

Male and female rats were immunized in the flanks with 200 μl of a 1:1 emulsion of ovalbumin (Sigma, St. Louis, MO) in complete Freund’s adjuvant (Difco/Becton Dickinson, Franklin Lakes, NJ)(25). After 7days, under isoflurane anesthesia, the rats were challenged with ovalbumin dissolved in saline in the pinna of one ear; the collateral ear was injected with saline (49). Rats received either a single subcutaneous injection of 0.1 mg/kg ShK-235 or P6N vehicle, a single oral gavage of 1×10^9 cfus LrS235 or LrGusA, oral gavage of 1 ml supernatant of LrS235, or a capsule filled with ShK-235 or gelatin immediately before the ear challenge. The DTH reaction was measured 24 hours post-challenge as the thickness of the ear using a spring-loaded micrometer (Mitutoyo, Japan) and ear inflammation was determined by comparing the ear thickness of the ovalbumin-challenged with the saline-challenged ear from each (49). The animals were euthanized after the ear measurements.

### Induction, monitoring, randomization, and treatment of rat collagen-induced arthritis

CIA was induced as described previously (46, 50). Briefly, female Lewis rats received a subcutaneous injection of 200□μl of a 1:1 emulsion of 2□mg/ml porcine type II collagen (20031, Chondrex, Redmond, WA) with incomplete Freund’s adjuvant at the base of the tail. After 7 days, rats were given a booster of 100□μl of collagen and adjuvant emulsion. Disease onset was defined as the development of at least one swollen or red paw joint. Clinical scores were determined daily by assigning 1 point for each swollen or red toe joint, 2 points for mildly swollen wrist or ankle joints, and 5 points for each severely swollen wrist or ankle, giving each rat a maximum possible score of 60. Upon disease onset, rats were treated every other day by the subcutaneous injection of P6N buffer vehicle or 0.1□mg/kg ShK-235, or the oral gavage of 1×10^9 cfus LrGusA or 1×10^9 cfus LrS235 daily. CIA is more severe in rats with early disease onset. To avoid biasing our results based on disease severity on the day each rat developed signs of disease and accounting for differences in the time between immunization and when a rat developed signs of illness, every rat that developed signs of disease on a given day was placed in a different treatment group to fill all groups in parallel.

### Histology and micro-CT

Healthy rats and rats from the CIA trials were euthanized after 21 days of treatment, and their hind paws were collected and fixed in 10% buffered formalin. One paw from each rat was imaged in the Optical Imaging & Vital Microscopy Core at Baylor College of Medicine by Micro-CT using a Bruker SkyScan 1272 Scanner set at 13 µm resolution with no filtering, no averaging, and a rotation step of 0.3. Raw images were analyzed with CTvox (Bruker, MA, US). The other hind paw was decalcified, embedded in paraffin, and sectioned by the Pathology & Histology Core at Baylor College of Medicine. Slides were stained with either hematoxylin and eosin or Safranin O/Fast green and imaged at 4x magnification on a Nikon Ci-L bright-field microscope (Nikon Inc. NY, US) in the Integrated Microscopy Core at Baylor College of Medicine. Scoring of the slides was completed by an investigator blinded to treatment groups using a comprehensive histological scoring system as described elsewhere (46, 51), in which cartilage degradation, cartilage erosion, angiogenesis, pannus formation, synovial hyperplasia, and synovial inflammation were evaluated by the following criteria: 0 = absent, 1 = mild, 2= moderate, 3 = severe. The data were presented as mean ± standard□deviation.

### Immunogenicity assays in rats with CIA

High-protein binding 96-well microplates (3855, ThermoFisher, MA, US) were coated overnight with 10 µg/ml of either porcine collagen II (20031, Chondrex, Woodinville, WI), ShK-235, or HsTX1[R14A], dissolved in PBS. Non-specific binding sites were blocked with PBS + 5% skimmed milk and washed with PBS + 0.05% Tween 20 before adding serum from the CIA rats, diluted in PBS. After washes, anti-rat IgG antibodies conjugated to horseradish peroxidase (1 µg/ml; Pierce catalog 31471, lot UA280036) were added to all the wells. One-step TMB-ELISA (34028, ThermoFisher, MA, US) was used to detect absorbance on a plate reader at 650 nm.

### Antibody neutralization assays

ShK-235 was pre-incubated with media supplemented with 10% of serum from the rats of the CIA trials or healthy, non-immunized and untreated rats. It was then added to the PBMCs loaded with CellTrace Violet (C34557, ThermoFisher, MA, US), and T cells proliferation was assessed as described above.

### Long-term immunogenicity assays in healthy rats

Healthy Lewis rats received a daily oral gavage of 1×10^9 cfus LrS235, 1×10^9 cfus induced LrGusA, or vehicle for 8 weeks. The rats were left untreated for 12 weeks and then were again treated daily for one week, reaching a total of 21 weeks. Blood was drawn at the 8 week and the 21 week time points and serum was tested for IgG against ShK-235 and HsTX1 [R14A] by ELISA, as described above.

### Statistical Analysis

Student’s T-test, one-way ANOVA, and two-way ANOVA without correction were used to determine whether differences among the groups were statistically significant (*P*<0.05). The CIA clinical scores analyses were completed using repeated measure one-way analysis of variance with Bonferroni post hoc test. Data were presented as mean ± SEM. All analyses were performed using GraphPad Prism.

## Acknowledgments

This project was funded in part by a pilot grant from the Alkek Center for Metagenomics and Microbiome Research at Baylor College of Medicine (to C.B. and J.M.H.) and by Bridge Funding from Baylor College of Medicine (to C.B.). The work was supported by the Cytometry & Cell Sorting core, the Pathology & Histology core, the Optical Imaging & Vital Microscopy Core, and the Integrated Microscopy Core funded in part by the Cancer Prevention and Research Institute of Texas (RP180672, RP150578, RP1806721, and RP170719), the National Institutes of Health (DK56338, CA125123, HG006348, and RR024574), the Dan L. Duncan Comprehensive Cancer Center, and the John S. Dunn Gulf Coast Consortium for Chemical Genomics.

