## Supplemental information for "A bioengineered probiotic for the oral delivery of a peptide Kv1.3 channel blocker to treat rheumatoid arthritis"

### Supplementary Information Text

### Materials and Methods

Supplemental Table

| Parameters | Healthy | LrGusA | ShK-235 | LrS235 |
|---|---|---|---|---|
| Synovial inflammation | <0.0001**** | 0.9853 | 0.0247* | 0.0097** |
| Synovial hyperplasia | 0.0002*** | 0.9731 | 0.0453* | 0.0007*** |
| Pannus formation | <0.0001**** | 0.9999 | 0.0219* | 0.0011** |
| Angiogenesis | <0.0001**** | 0.9947 | <0.0001**** | <0.0001**** |
| Cartilage erosion | <0.0001**** | >0.9999 | 0.0123* | 0.0013** |
| Cartilagedegradation | 0.0002**** | >0.9999 | 0.0406* | 0.0065** |

**Table S1**. P values for 2-way ANOVA analysis, compared with vehicle-treated CIA rats. 2-way ANOVA analysis was performed, followed by Tukey’s multiple comparisons test, with individual variances computed for each comparison. *P < 0.05; **P < 0.01, ***P< 0.001, ****P< 0.0001.

Supplemental Figures

**A**

ShK RSCIDTIPKSRCTAFQCKHSMKYRLSFCRKTCGTC
ShK-186 **pTyr-AEEA-**RSCIDTIPKSRCTAFQCKHSMKYRLSFCRKTCGTC
ShK-235 RSCIDTIPKSRCTAF**K**CKHS**I**KYRLSFCRKTCGTC**A**

**B**

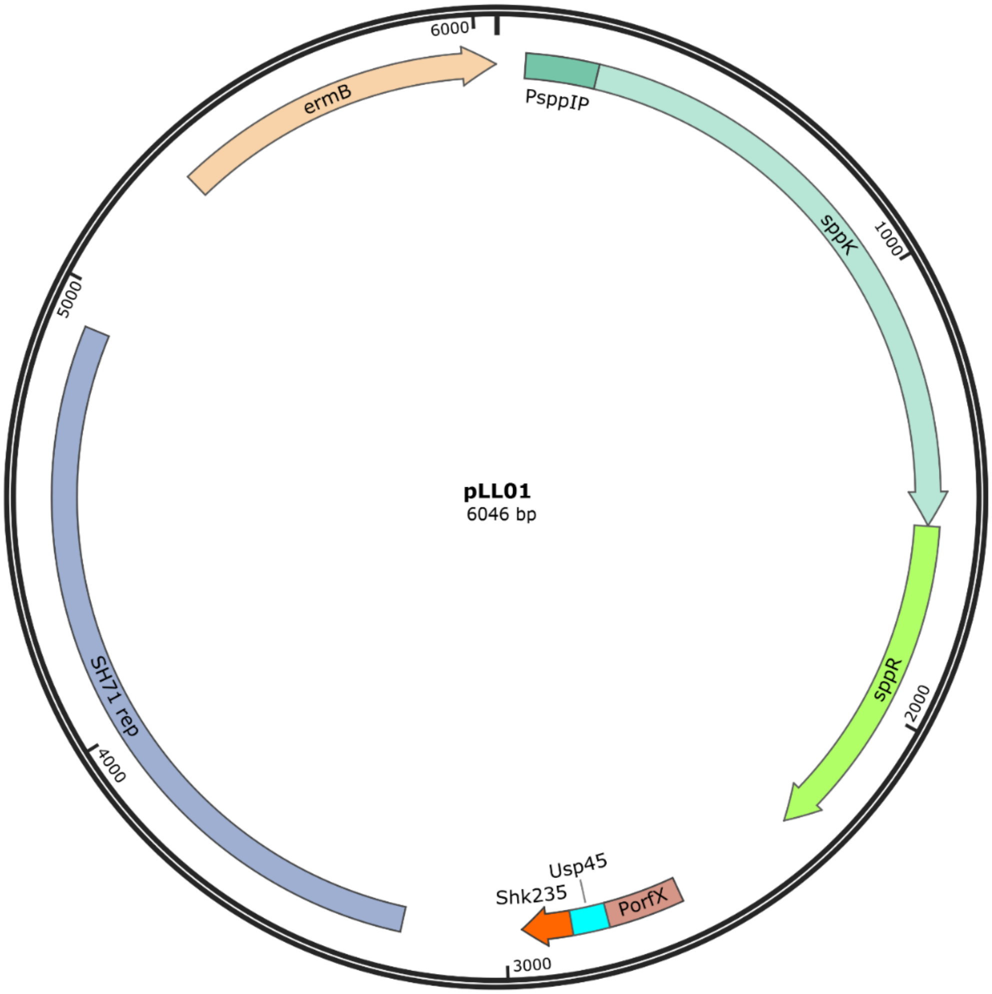


**Figure S1**. **Design of the inducible expression system. A,** Amino acid sequences of ShK, its synthetic analog ShK-186, and its recombinant analog ShK-235. Differences between sequences are shown in bold and red. ShK is the 37 amino acid peptide originally isolated from the venom of the sea anemone *Stichodactyla helianthus* (1). ShK-186 contains a pTyr attached to the peptide's N-terminus via a 9-carbon atom linker (AEEA) that precludes its recombinant production. ShK-235 differs from ShK by a Q16K substitution, an I21M substitution, and the addition of an Ala to the C-terminus. **B,** Codon optimized ShK235 with signal peptide Usp45 was ligated into *Nco*I-*Xho*I digested pSIP411 resulting in secretion expression plasmid pLL01.

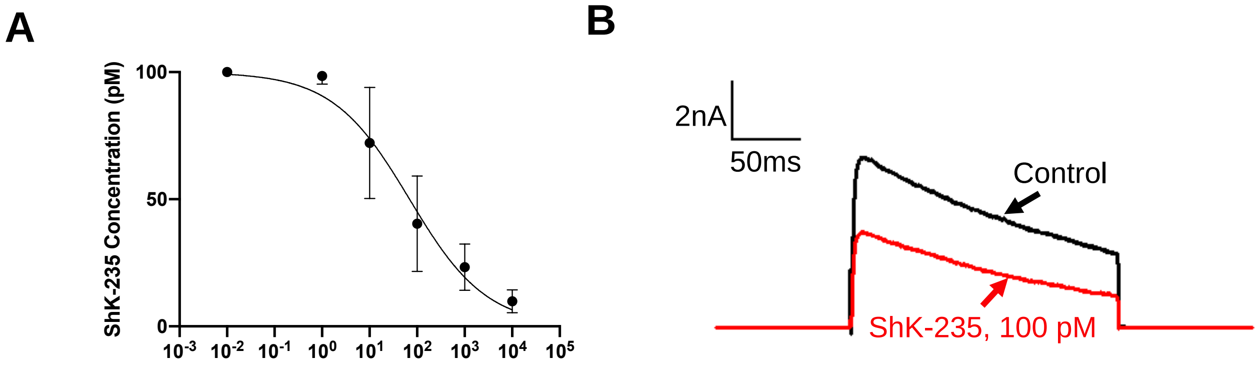


**Figure S2. Dose-response of synthetic ShK-235 block of Kv1.3.**
A, Dose-response of mKv1.3 block by synthetic ShK-235. N = 3 cells per concentration. IC50 = 69.31 ± 24.0 pM. B, Representative traces of the block of mKv1.3 currents by 100 pM synthetic ShK-235.

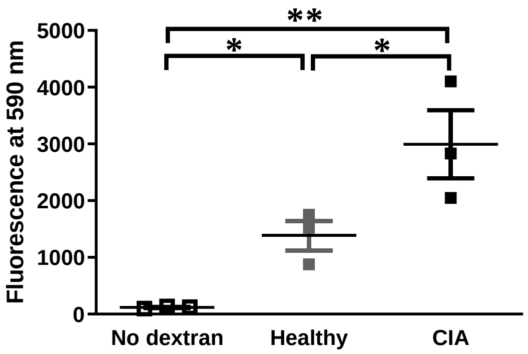


**Figure S3. Intestinal permeability of healthy rats and rat with CIA to 4 kDa dextran.**
Healthy rats (□, ■) and rats at the onset of CIA (■) received an oral bolus of 60 mg/kg FITC-labeled 4 kDa dextran (filled squares) or saline (open squares). A sample of blood was drawn 6 hrs later and serum levels of FITC fluorescence were measured at 590 nm with a fluorescence plate reader. N = 3 rats per group. *p<0.05, **p<0.01.

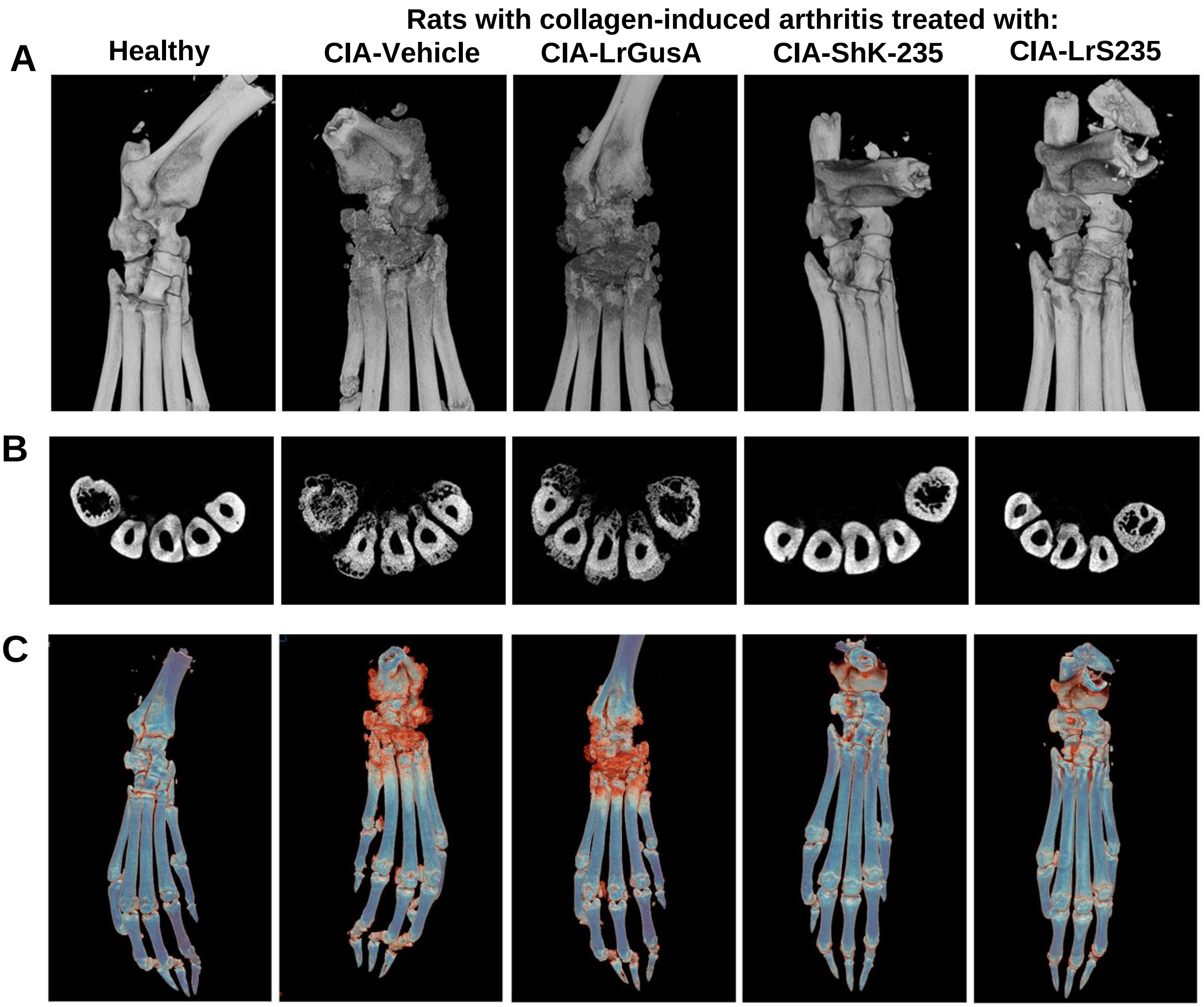


**Figure S4. Microfocal computed tomography (Micro-CT) validation of the therapeutic effects of LrS235.** Micro-CT scan validates the therapeutic effects of engineered probiotic LrS235 on bone destruction in rats with collagen-induced arthritis (CIA). Rats were injected with vehicle (P6N buffer), synthetic ShK-235 (100 μg/kg) every other day, or orally administered with LrGusA ($1x10^9$ CFU) or LrS235 ($1x10^9$ CFU) daily for 21 days from the first day of the onset of the clinical symptoms of arthritis. Micro-CT scan was performed to assess bone damage at the end of the experiments. (A) 3D reconstructed bones of ankle in different groups. (B) Cross-section at the level of metatarsals. (C) Pseudocolor of micro-CT data from fig. S4 A. Red color corresponds to low density pixels, indicating area of bone erosion and cartilage damge, blue color corresponds to high density pixels, indicating high density bone.

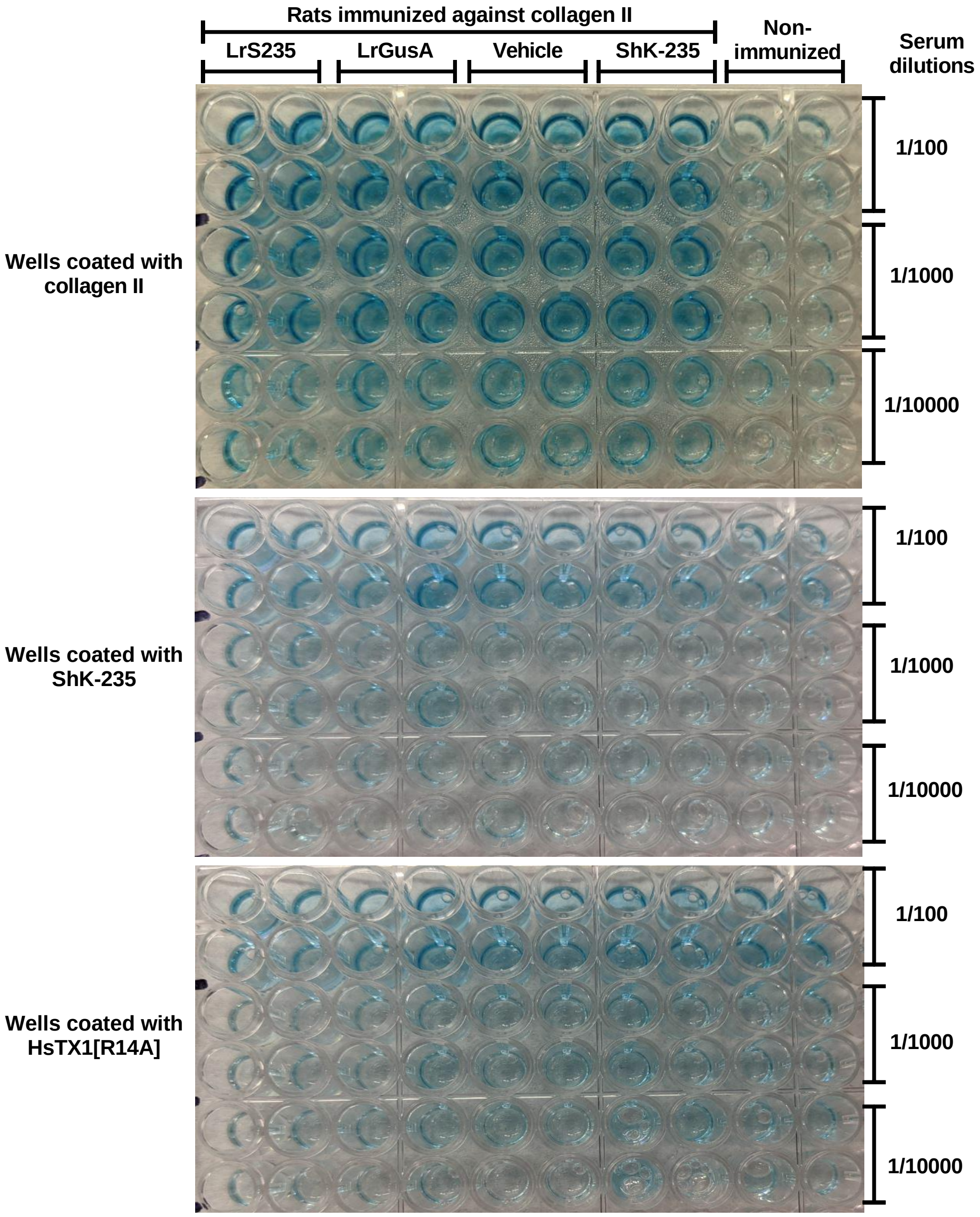


**Figure S5. Representative photos of the ELISA plates from figure 4A.** ELISA plates were coated with 10 µg/ml of either collagen II, ShK-235, or the Kv1.3 blocker HsTX1[R14A].

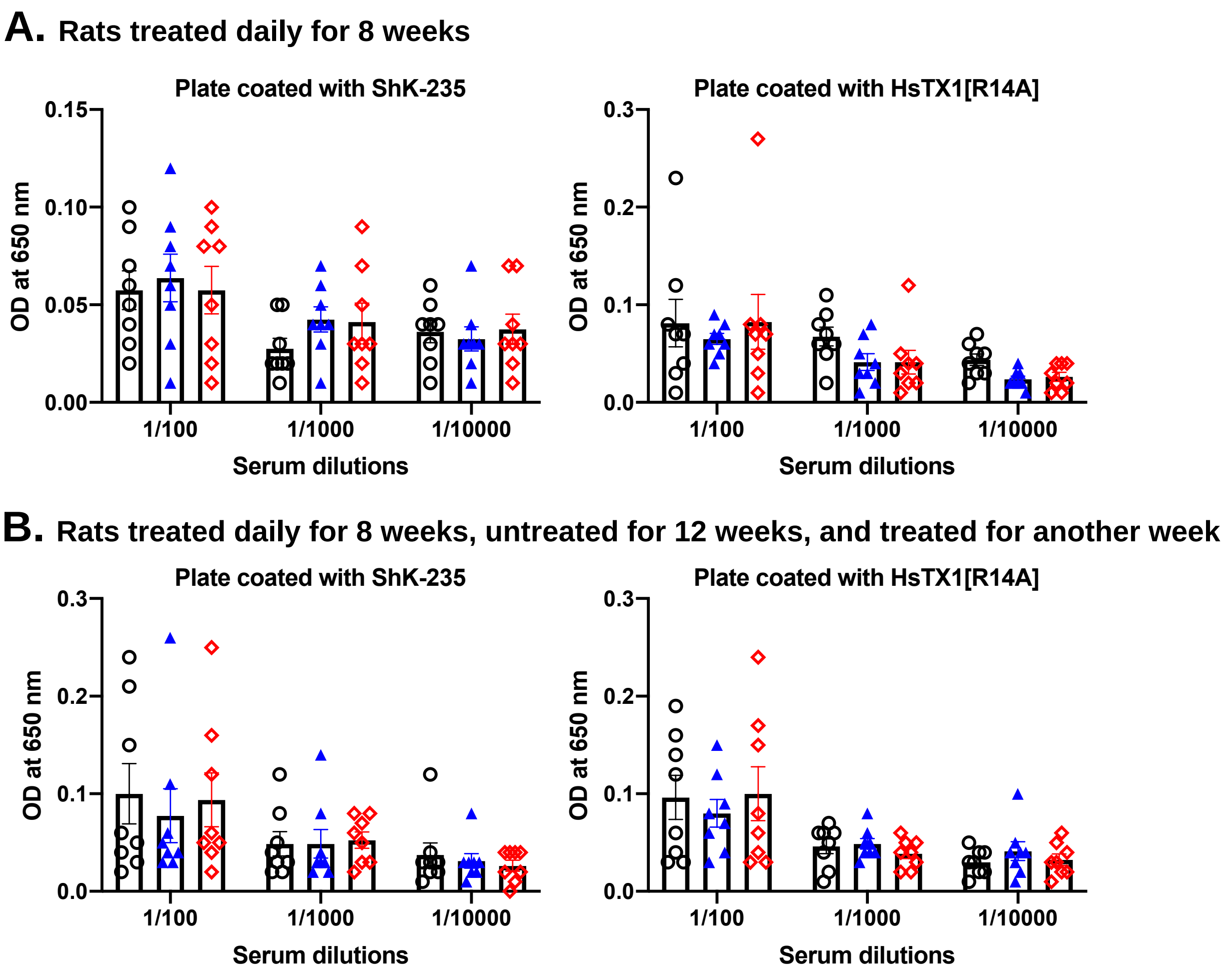


**Figure S7. Lack of antibodies against ShK-235 in rats after 8 weeks of treatment.** ELISA plates were coated with 10 µg/ml of ShK-235 or the Kv1.3 blocker HsTX1[R14A] with no sequence or structure homology to ShK-235, before incubation with serum from the healthy rats treated daily for 8 weeks (A) or treated daily for weeks, untreated for 12, and treated again daily for 1 week (B) with vehicle (○), LrGusA (▲), or LrS235 (◇). Serum was diluted 1:100, 1:1000, and 1:100000, and bound IgG were detected using anti-rat IgG-HRP and TMB substrate at 650 nm. N = 8 rats per group (4 males, 4 females).
